# Repeatability and reproducibility assessment in a large-scale population-based microbiota study: case study on human milk microbiota

**DOI:** 10.1101/2020.04.20.052035

**Authors:** Shirin Moossavi, Kelsey Fehr, Theo J. Moraes, Ehsan Khafipour, Meghan B. Azad

## Abstract

**Background:** Quality control including assessment of batch variabilities and confirmation of repeatability and reproducibility are integral component of high throughput omics studies including microbiome research. Batch effects can mask true biological results and/or result in irreproducible conclusions and interpretations. Low biomass samples in microbiome research are prone to reagent contamination; yet, quality control procedures for low biomass samples in large-scale microbiome studies are not well established.

**Results:** In this study we have proposed a framework for an in-depth step-by-step approach to address this gap. The framework consists of three independent stages: 1) verification of sequencing accuracy by assessing technical repeatability and reproducibility of the results using mock communities and biological controls; 2) contaminant removal and batch variability correction by applying a two-tier strategy using statistical algorithms (e.g. *decontam*) followed by comparison of the data structure between batches; and 3) corroborating the repeatability and reproducibility of microbiome composition and downstream statistical analysis. Using this approach on the milk microbiota data from the CHILD Cohort generated in two batches (extracted and sequenced in 2016 and 2019), we were able to identify potential reagent contaminants that were missed with standard algorithms, and substantially reduce contaminant-induced batch variability. Additionally, we confirmed the repeatability and reproducibility of our reslults in each batch before merging them for downstream analysis.

**Conclusion:** This study provides important insight to advance quality control efforts in low biomass microbiome research. Within-study quality control that takes advantage of the data structure (*i.e.* differential prevalence of contaminants between batches) would enhance the overall reliability and reproducibility of research in this field.

## Background

Quality control of microbiome studies has been an integral component of pioneering projects including the Human Microbiome Project [1]. The Microbiome Quality Control Project (MQCP) focused on identifying sources of variability in 16S rRNA gene microbiota profiling across different laboratories, but batch-to-batch variability was not assessed [2]. As microbiome studies expand in sample size, we are facing the additional challenge of batch-to-batch variability in large-scale population-based studies. Additionally, repeatability and reproducibility of results are often unaddressed. Unlike other high-throughput methods such as transcriptomics and metabolomics [3, 4], these concepts are not well developed for microbiome studies.

“Batch effects are sub-groups of measurements that have qualitatively different behaviour across conditions and are unrelated to the biological or scientific variables in a study” [4]. Batch effects can mask true biological results and/or result in irreproducible conclusions and interpretations [4]. Potential sources of batch effects in microbiome research include heterogeneity in all aspects from sample collection to library preparation and bioinformatics processing [1] leading to technical variability. Reagent contaminants pose a major challenge in microbiome profiling of low biomass samples such as milk [5, 6]; and could be an important source of non-technical batch variability even when all procedures are identical.

Repeatability is defined as obtaining the same results after re-running the same process on the same set of samples, while reproducibility refers to the ability to obtain similar results on a different set of samples [7]. Assessing repeatability and reproducibility is among the cornerstones of good scientific conduct and is being adopted in many areas of high-throughput experiments such as clinical genomics [8]. Studies have assessed the reproducibility of the microbiome profile as part of MQCP [2]. However, repeatability and reproducibility of results are not commonly assessed between batches. This process is important when combining results from multiple batches in large-scale microbiome projects. Therefore, the objective of this study was to perform extensive quality control and establish good practices using milk microbiome data generated in two batches (extracted and sequenced in 2016 and 2019). Additionally, we assessed and mitigated batch variability and examined repeatability and reproducibility in this dataset.

## Results

We studied a subset of 1,194 mother-infant dyads in the CHILD Cohort Study [9]. Milk microbiota from a representative subset of 428 mothers was previously profiled in 2016 (Batch 1) [10]. An additional set of 766 samples enriched in infant atopy and asthma was profiled in 2019 (Batch 2). Experimental and bioinformatics procedures were identical for the two batches with the exception of DNA extraction kit lots. Some participant characteristics varied significantly between the batches (e.g. season of birth differed and atopy/asthma were purposefully enriched in Batch 2; **Table S1**) and thus some degree of true biological variability between batches was anticipated in the milk microbiota composition (Supplementary Methods).

### Technical reproducibility

Technical reproducibility of library preparation and sequencing was confirmed on a mock community consisting of DNA extracted from 8 bacterial species (Zymo Research, USA) and biological controls (comprising of 9 Batch 1 samples re-sequenced in Batch 2; Figure 1A&B). Substantial inter-individual variability was observed, as expected. However, the composition remained consistent between batches within each individual (Figure 1B).

**Figure 1.**
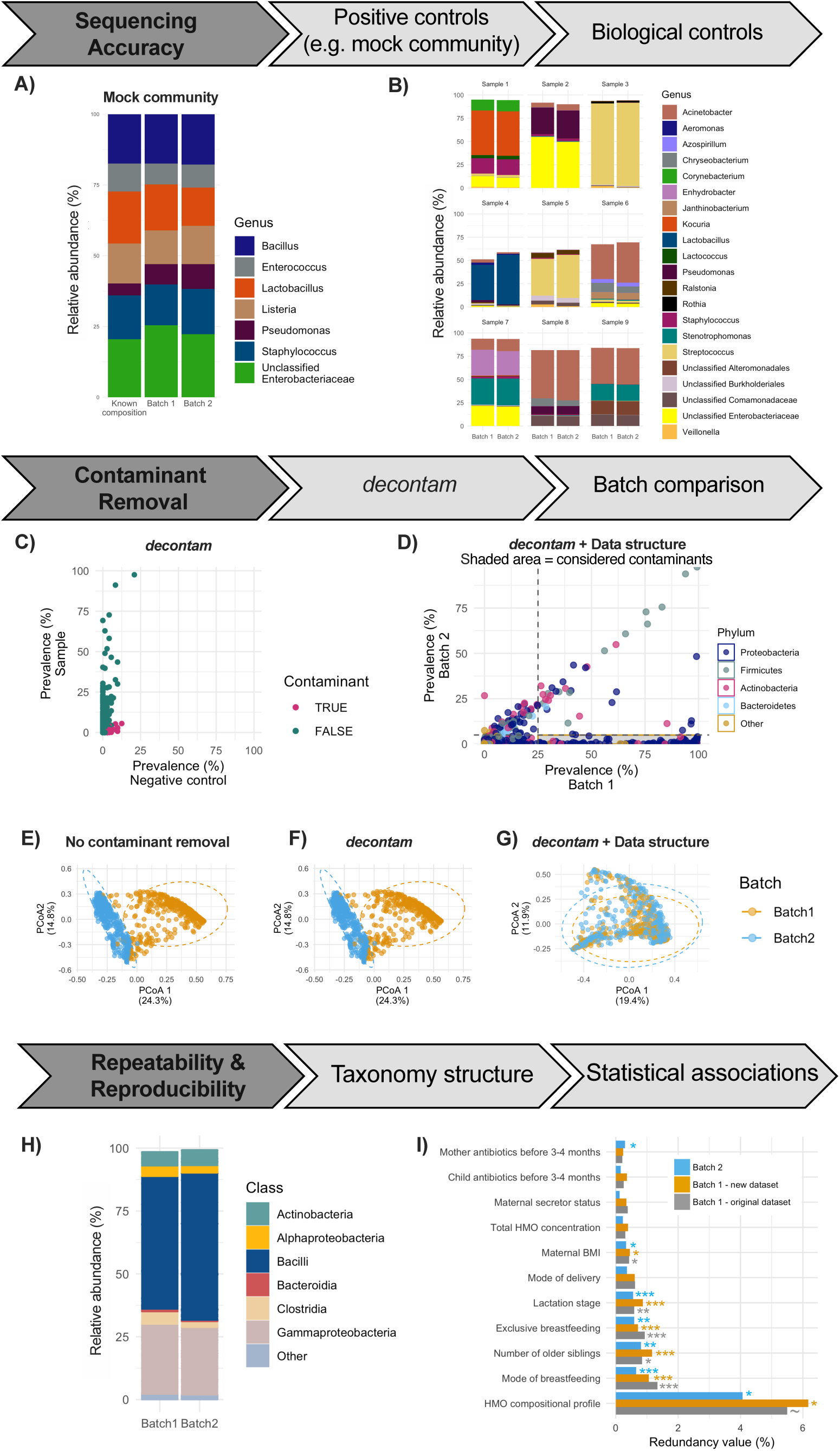
Multi-layer quality control approach to assess reproducibility and repeatability in a large-scale milk microbiota study. Sequencing technical accuracy was verified on A) mock community and B) biological controls comprising of 9 samples originally processed in Batch 1 that were re-sequenced in Batch 2. Next, potential reagent contaminants were identified using a two-tier strategy including C) *decontam* package and D) using the data structure by comparing ASV prevalence between batches. ASVs with more than 25% prevalence in one batch and less than 5% prevalence in the other batch were considered contaminants. ASVs identified in either of methods C or D were considered reagent contaminants and removed. Batch variability was assessed E) prior to contaminant removal, F) after *decontam* [12], and again G) after considering the data structure *i.e.* taxa prevalence between the batches [13]. The two-tier strategy eliminated the prominent separation of the samples on the PCoA plot assessed on the Bray-Curtis dissimilarity. Subsequently, repeatability and reproducibility of the results were assessed. H) Comparison of milk microbiota taxonomy between the two batches. I) Comparison of the statistical associations of determinants of the milk microbiota composition using redundancy analysis as previously described [10].

### Two-tier strategy using the decontam algorithm and milk microbiota data structure to identify reagent contaminants

As milk is a low biomass sample, reagent contaminants could plausibly be present in the sequencing output of samples [11] and thus major differences between the batches could be due to different profiles of the reagent contaminants. A two-tier strategy was used to identify potential reagent contaminants followed by assessing the milk microbiota variability between batches. First, potential reagent contaminants (N=256 ASVs) were identified and removed using the *decontam* package based on either the frequency of the ASV in negative controls or the negative correlation with DNA concentration [12] (Figure 1C). The negative controls included were extraction negative control for Batch 2 and sequencing negative control for both batches. Second, we followed the recommendation of de Goffau et al. based on the premise that “within-batch consistency of the reagent contamination profile and between batch variation of such profiles are two of the most powerful tools that can be used to recognize reagent contamination” [13]. In other words, we expect correlation in true signals between different batches while potential contaminants will not be correlated [13]. We took advantage of the data structure of each batch to identify additional reagent contaminants that were not identified by the *decontam* algorithm. While there was a high degree of correlation in the prevalence of the majority of ASVs between the two batches, 113 ASVs were present in over 25% of samples in Batch 1 and less than 5% in Batch 2 (Figure 1D); these ASVs were considered contaminants and removed.

### Reagent contaminants as the major source of batch variability

Despite the technical reproducibility, which was confirmed on mock community and biological controls (Figure 1A&B), preliminary comparisons between batches revealed differences in beta diversity of milk microbiota (Figure 1E). This difference remained after applying *decontam*, which identified reagent contaminants in both batches (Figure 1F). However, removing the additional potential contaminants from Batch 1 identified through comparison of the data structure between batches eliminated the differences in milk microbiota composition between the batches (Figure 1G).

### Repeatability and reproducibility assessment

Next we assessed the repeatability and reproducibility of the milk microbiota composition taxonomy and statistical associations with its determinants. The composition of the core ASVs (especially those suggested to be common reagent contaminants: *Comamonadaceae*, *Rhodospirillaceae*, and Burkholderiales) was affected by the updated pre-processing and contaminant removal. As a result, the core microbiota as previously defined (ASVs present in at least 95% of samples with at least 1% mean relative abundance) [10] was not repeatable or reproducible, underscoring the challenges of defining core taxa in low biomass samples. However, the taxonomic structures of the most abundant taxa were consistent between the two batches (Figure 1H).

Next, the robustness of the associations with determinants of the milk microbiota composition was assessed. We had previously performed redundancy analysis (RDA) on Batch 1 to identify factors associated with the overall composition of milk microbiota [10]. We repeated the same RDA analysis on the re-processed Batch 1 and the new Batch 2 milk microbiota composition (Figure 1I). We confirmed repeatability of the results within Batch 1, despite removing several ASVs during the updated pre-processing. Additionally, most of the associations including mode of breastfeeding were reproduced in Batch 2 (Figure 1I). Based on these results, we felt confident to merge the two datasets for our ongoing research.

## Discussion

Rigorous quality control and assessment of repeatability and reproducibility of results are infrequently reported for microbiome studies. Here, we have proposed a framework for an in-depth step-by-step approach to a comprehensive quality control assessment of low biomass microbiome. The framework consists of three independent stages: 1) verification of sequencing accuracy by assessing technical repeatability and reproducibility of the results using mock communities and biological controls; 2) contaminant removal and batch variability correction by applying a two-tier strategy using statistical algorithms (e.g. *decontam*) followed by comparison of the data structure between batches; and 3) corroborating the repeatability and reproducibility of microbiome composition and downstream statistical analysis.

Batch variability can be minimized by adhering to standardised protocols and using consumables of the same lot; however, the latter may be impractical in longitudinal studies or when samples are analyzed over a prolonged period of time, as in our study. Methods to minimise batch variability post-analysis have been developed based on various normalisation approaches, which generally assume that the batch variability is due to random technical variations [14, 15]. However, these global normalisation approaches (e.g. quintile normalisation) cannot eliminate the batch effect in variables that are differentially impacted in different batches [4] - for example, if the batch effect is due to differing reagent contaminant profiles instead of random technical variabilities.

### Conclusion

Our study highlights the importance of reagent contaminants as a potential source of batch variability in low biomass samples [16]. Guidelines have been developed to minimise the influence of contaminants in low biomass samples [17, 18]. However, these do not extend to contaminant-related batch variation. Our results indicate that conducting a comprehensive quality control assessment when profiling the microbiome of milk and other low biomass samples would ensure more robust, generalizable, and reproducible results. Specifically, we recommend within-study quality control that takes advantage of the data structure (*i.e.* differential prevalence of contaminants between batches) to enhance the overall reliability and reproducibility of research in this field [19].

## Methods

### Study design

We studied a subset of 1,194 mother-infant dyads in the Canadian Healthy Infant Longitudinal Development (CHILD) birth cohort, designed to study the developmental origins of pediatric asthma and allergy [9]. Women with singleton pregnancies were enrolled between 2008 and 2012 and remained eligible if they delivered a healthy infant >35 weeks gestation (n=3455). Milk microbiota from a representative subset of 428 mothers was previously profiled (2016; Batch 1) [20]. An additional set of 766 samples enriched in infant atopy and asthma was included in this study (2019; Batch 2). Participants gave written informed consent in accordance with the Declaration of Helsinki. The protocol was approved by the Human Research Ethics Boards at McMaster University, the Hospital for Sick Children, and the Universities of Manitoba, Alberta, and British Columbia.

### Sample collection and microbiota analysis

Each mother provided one sample of milk collected during a 24-hour period at 4 months postpartum [mean (SD) 17 (5) weeks postpartum] as previously described (Moossavi et al. submitted). Batch 2 samples were processed similar to Batch 1 as previously described [10]. Briefly, genomic DNA was extracted from 1 ml breastmilk using Quick-DNA Fungal/Bacterial extraction kit following the manufacturer’s instructions (Zymo Research, USA). Extraction kits were purchased separately for Batch 1 and Batch 2. Samples were sequenced following amplification of the V4 hypervariable region of the 16S rRNA gene with modified F515/R806 primers [21] on a MiSeq platform (Illumina, San Diego, CA, USA) in 2016 (Batch 1) and 2019 (Batch 2). Sterile DNA-free water was used as negative controls in in the DNA extraction (only Batch 2) and sequencing library preparation (Batch 1&2). A mock community consisting of DNA extracted from of 8 bacterial species with known theoretical relative abundances (Zymo Research, USA) was included as positive control in sequencing library preparation. Genomic DNA of 9 milk samples previously sequenced in Batch 1 were also included in sequencing library preparation of Batch 2 as biological controls.

Overlapping paired-end reads were processed with DADA2 pipeline [22] using the open-source software QIIME 2 v.2018.6 (https://qiime2.org) [23]. Unique amplicon sequence variants (ASVs) were assigned a taxonomy and aligned to the 2013 release of the Greengenes reference database at 99% sequence similarity [24]. Demultiplexed sequencing data was deposited into the Sequence Read Archive (SRA) of NCBI and can be accessed via accession numbers PRJNA481046 and PRJNA597997.

### Microbiome pre-processing and reagent contaminant identification

Data analysis was conducted in R [25]. Initial preprocessing of the ASV table was conducted using the Phyloseq package [26]. Potential reagent contaminants were identified using *decontam* package based on either the frequency of the ASV in the negative control or the negative correlation with DNA concentration [12]. Additional potential contaminants were identified by comparing the prevalence of ASVs between batches as previously suggested [13]. ASVs with more than 25% prevalence in one batch and less than 5% prevalence in the other batch were also considered contaminants and removed. Subsequently, ASVs only present in the mock community or negative controls (n=1,570), unassigned ASVs, and ASVs belonging to the phylum Cyanobacteria, family of mitochondria and class of chloroplast (n=780) were removed. Samples were rarefied to the minimum of 8,000 sequencing reads per sample resulting in 877 samples and 9,985 remaining ASVs. ASVs with less than 60 reads across the entire dataset were also removed, resulting in 1,122 remaining ASVs. The number of sequencing reads per sample was then relativized to a total sum of 8,000 for downstream analyses.

### Data quality control assessment

Technical reproducibility was assessed by agreements in taxonomic structure of biological controls, mock community, and milk microbiota between batches. The batch effect was assessed on the overall milk microbiota composition using Bray-Curtis dissimilarity and visualised in PCoA plots. Repeatability was verified by examining the associations of maternal, infant, and early life factors with milk microbiota using redundancy analysis (RDA) in the original Batch 1 [10], new re-processed Batch 1, and Batch 2.

## List of abbreviation

MQCP: Microbiome Quality Control Project
ASV: Amplicon sequencing variant
RDA: Redundancy analysis
PCoA: Principal coordinate analysis
CHILD: Canadian Healthy Infant Longitudinal Development

## Declarations

### Ethics approval and consent to participate

Written informed consent was obtained from the participants. The original study was approved by the Human Research Ethics Boards at McMaster University, the Hospital for Sick Children, and the Universities of Manitoba, Alberta, and British Columbia.

### Consent for publication

Not applicable

### Availability of data and material

The datasets generated and/or analysed during the current study are available in the Sequence Read Archive of NCBI repository (accession number PRJNA481046 and PRJNA597997).

### Authors’ contributions

Conceptualization, SM, MBA; Methodology, SM; Investigation, SM, KF; Writing – Original Draft, SM, MBA; Writing – Review & Editing, KF, TJM, EK. All authors read and approved the final manuscript.

## Acknowledgements

We are grateful to all the families who took part in this study, and the whole CHILD team, which includes interviewers, nurses, computer and laboratory technicians, clerical workers, research scientists, volunteers, managers, and receptionists. We thank Padmaja Subbarao (University of Toronto) Director of the CHILD study, Malcolm R. Sears (McMaster University) Founding Director, and CHILD site directors: Stuart E. Turvey (University of British Columbia), Allan B. Becker (University of Manitoba), and Piushkumar J. Mandhane (University of Alberta) for their leadership support; Diana Lefebvre (McMaster University) for managing the CHILD database; Leah Stiemsma for creating the antibiotic variables; and Lorena Vehling for creating the breastfeeding variables..

